# Modeling American Kestrel Decline Using Spatiotemporal Subsampling to Improve eBird Data Reliability

**DOI:** 10.1101/2024.10.31.621376

**Authors:** Archith Sharma, James C. Bednarz, Junhyeon Kwon

## Abstract

American Kestrel (Falco sparverius) numbers have plummeted over 40% since 1980, despite many other raptor species, including the closely related Merlin (Falco columbarius) rebounding in the same period, post the ban on the use of DDT. One plausible reason for this decline is the increase in Cooper’s Hawks (Astur cooperii), a predatory species. To model the impact of Cooper’s Hawk on American Kestrels, one can use citizen science data such as eBird. However, eBird contains a heavy sampling bias towards recent data in large cities as it has gained popularity over time. A novel framework testing different spatiotemporal subsampling resolutions was employed in seeing how to make the data more reliable and to enjoy a finer time resolution compared to the Breeding Bird Survey (BBS) standard. The ecological relationship between two species is then modeled using a time-series framework that accounts for both current and past interactions, with the model order determined based on criteria such as BIC. The creation and analysis of synthetic data are also tested to verify that our framework reflects the ground truth of the bird abundance trend. Immediate forecast results based on the eBird data suggests a decline of American Kestrels at the current pace.

## 1 Introduction

The American Kestrel (*Falco sparverius*), seen in Figure 1 is North America’s smallest and most abundant falcon. While Kestrels are about the same size as a mourning dove, they are known to be vicious predators. Kestrels are regular breeding inhabitants in most of North America. They prefer open, grassy habitat in search of their prey, mostly insects.

**Figure 1:**
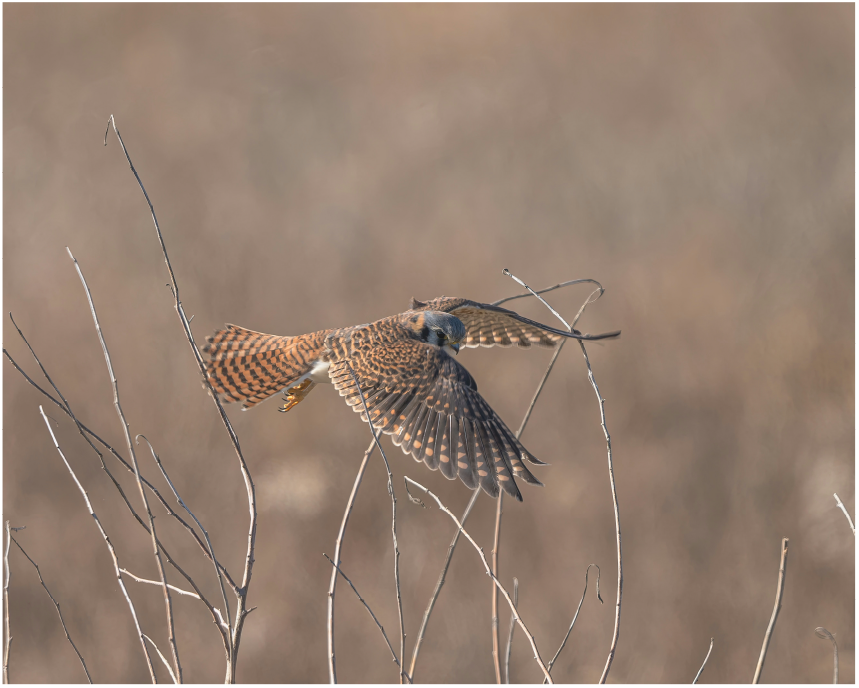
Female American Kestrel

In total, North America has lost an estimated 2 million Kestrels since 1970, despite the ban on DDT, according to the Peregrine Fund, a leading organization in raptor conservation. Conservancy agencies and past research point to the decline being attributed to various potential factors, including predation by Cooper’s Hawks (*Astur cooperii*), habitat change, and pesticide use (Sullivan et al., 2009).

This paper focuses on the hypothesis that the predation by Cooper’s Hawks has contributed to the decline of Kestrel populations. This ecological relationship between two species can be better explained by accounting for temporal dependence. For this purpose, we first need to understand the characteristics of the observation data for these species from two primary sources: eBird and the Breeding Bird Survey (BBS). eBird is a public repository of bird observations maintained by the Cornell Lab of Ornithology (Sullivan et al., 2009). The dataset contains records of bird sightings reported by the general public. Birders enter where they birded (location), for how long, and what birds they saw. This data is uploaded to the eBird database, where automated systems and regional reviewers maintain data quality. eBird has grown tremendously over time, with over 1 billion observations recorded as of 2024. As a result, eBird data provides extensive spatiotemporal coverage but includes sampling biases. On the other hand, BBS is a long-term dataset collected through annual surveys of bird populations during the breeding season (June) across North America maintained by the United States Geological Survey and the Canadian Wildlife Service (Ziolkowski Jr. et al., 2023; Sauer et al., 2017). The survey consists of thousands of fixed routes over uniformly distributed regions across the United States and Canada as shown in Figure 2. As BBS is regarded as a standardized and more reliable dataset collected through systematic protocols, this research uses it to verify the quality of debiased eBird data.

**Figure 2:**
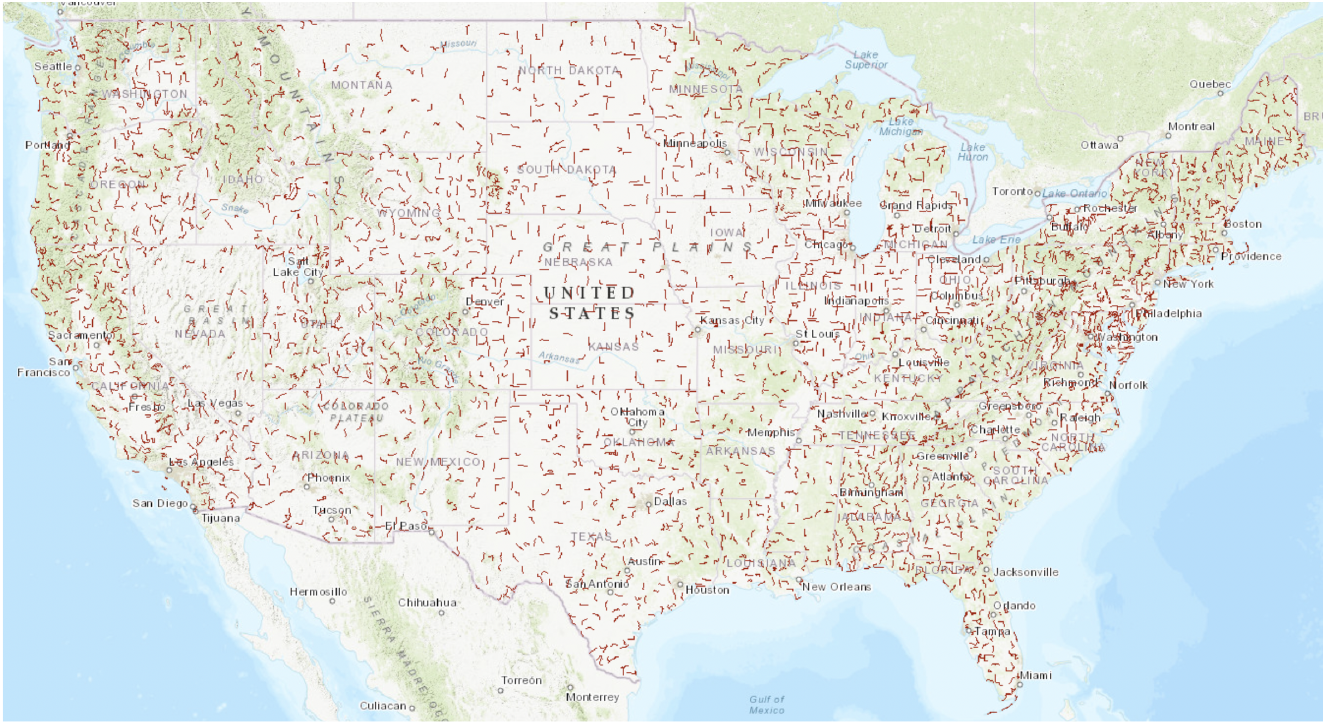
BBS Annual Survey Routes

To address inherent biases in eBird data while benefiting from its finer temporal resolution, we propose a two-fold framework consisting of i) subsampling-based debiasing to mitigate spatiotemporal sampling issues and ii) time series analysis that captures both current and past interactions between Kestrels and Cooper’s Hawks. Subsampling mitigates the overrepresentation of regions near large cities by randomly selecting one checklist from each spatiotemporal grid cell, following the approach of Johnston et al. (2021). The grid cell size is determined separately for each species based on the degree of similarity between the eBird dataset and the BBS dataset. The time series analysis method used in this paper incorporates the abundance variables of both species, as well as their temporal lags, to account for lag dependency in their interactions. For instance, the abundance of Kestrels of a given year may be influenced by their own abundance or that of Cooper’s Hawks in the previous period. This approach provides a more comprehensive understanding of the dynamic ecological relationship between the two species over time.

The paper is organized as follows. Section 2 presents our methods for debiasing eBird data and their application to estimate and forecast abundance trends for focal bird species using real-world data from the Atlantic Waterfowl Flyway. Section 3 details the simulation study designed to verify the framework’s ability to capture trend dynamics in the presence of spatiotemporal sampling bias in synthetic data. Finally, Section 4 concludes with an interpretation of the model fitting and forecast results, discussing the implications, contributions, and limitations of our approach and potential directions for future research.

## 2 Methods and Application

This section presents a framework designed to explain the ecological relationship between American Kestrels and Cooper’s Hawks. Using real-world data from the Atlantic Waterfowl Flyway, we illustrate how the proposed framework addresses sampling bias of eBird data. Subsequently, the framework is applied to uncover interspecies relationships and generate abundance forecasts, as depicted in Figure 3.

**Figure 3:**
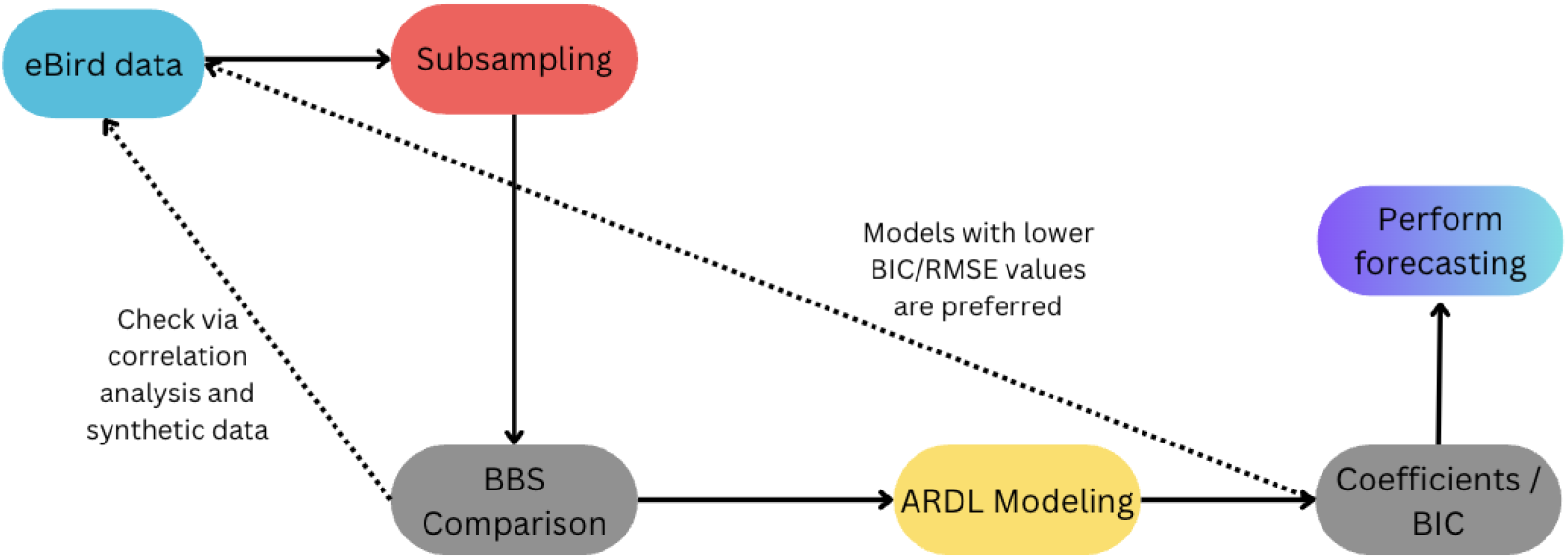
Methods for eBird time series

### 2.1 Data Specification

The proposed method is applied to species abundance data collected in the Atlantic Waterfowl Flyway in the United States for the time period between 1976 and 2022. The Atlantic Waterfowl Flyway consists of the Eastern United States, which is the region where Kestrels have seen the most noticeable decline in the last 40 years (Bednarz et al., 1990). Even though eBird’s recordkeeping began in 2002, data prior to that exists in the eBird archives since many users have uploaded prior checklists before eBird was created. This extra data creates a historical context for current population trends.

### 2.2 Spatiotemporal Subsampling

Despite the measures in place to ensure eBird data quality, biases are present in eBird data, most notably spatial and temporal bias. Users sample near their homes or in accessible areas (Luck et al., 2003). eBird has also had an exponential growth in the number of users since its launch, leading to a strong bias towards recent data. The full plot of eBird data (at different time intervals) is present in Figure 4, and we can observe that …

**Figure 4:**
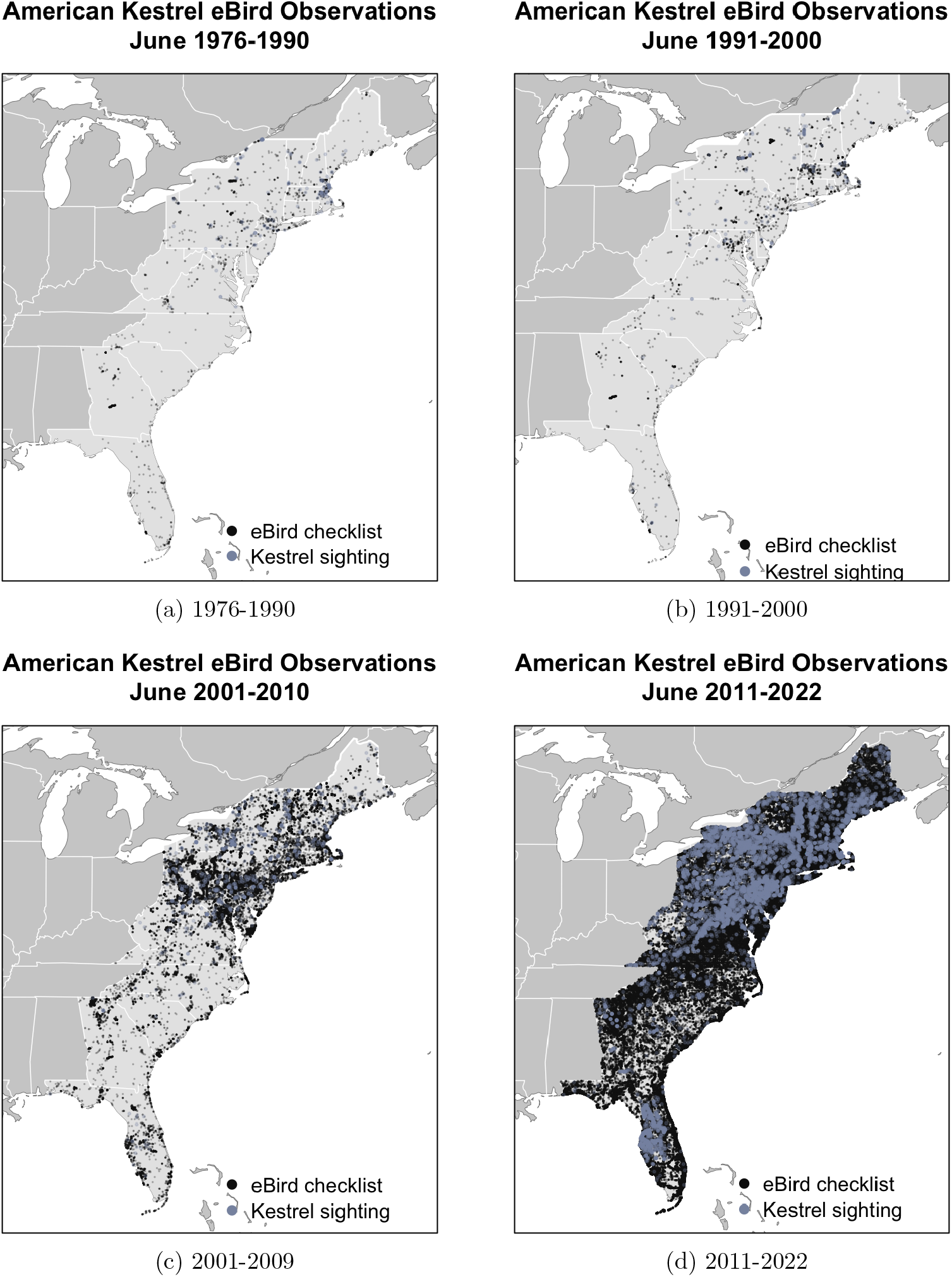
Spatiotemporal sampling bias of eBird data.

To address spatial overrepresentation, we subsample the eBird checklists as in Johnston et al. (2021) by selecting one checklist per spatial grid cell. Temporal bias from recent years is mitigated by normalizing bird observation counts by the number of checklists in eBird data or by the number of routes in the BBS dataset. This subsampling approach ensures a more balanced representation of both spatial and temporal dimensions, improving the reliability of the data for subsequent analyses.

To explain the subsampling approach more in detail, … Creating equal area grids in the study region and sampling a given amount per box for each year address both of these biases, as shown in Figure 5.

**Figure 5:**
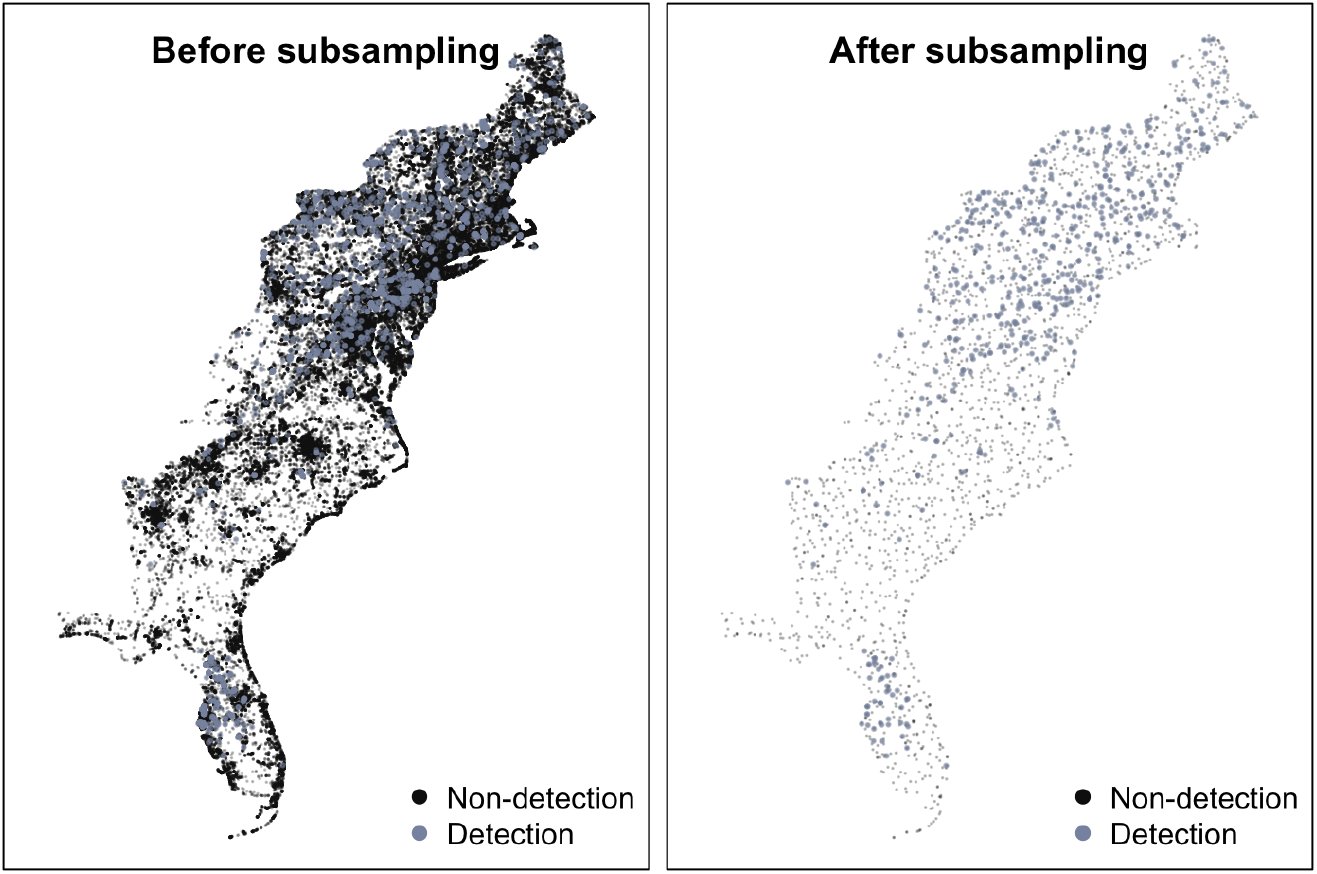
30 km subsampled grid

The spatial grid size for subsampling (referred to as the subsampling resolution) is optimized by assessing how closely the results approximate the BBS data during the breeding month (June). For each species, we perform subsampling of the eBird data at varying spatial resolutions (1–30 km), repeating the process 200 times for each resolution. For each resolution, the median of the subsampled annual abundance is calculated. The optimal resolution is identified as the one that achieves the highest correlation with the corresponding BBS data.

As shown in Figure 8, for Kestrels, the 1 km resolution most closely approximated the BBS data. In contrast, for Cooper’s Hawks, the optimal resolution was 29 km, likely reflecting differences in the spatial scales at which the species are observed. Figure 7 illustrates the abundance trends for each species based on eBird data at their respective optimal sub- sampling resolutions. The subsampling process noticeably improved the alignment of eBird trends with those from the BBS dataset in Figure, providing a more reliable representation of species abundance over time. This optimized subsampling approach is then applied to the entire monthly eBird dataset, providing the basis for fitting a time series model in the subsequent analysis.

**Figure 6:**
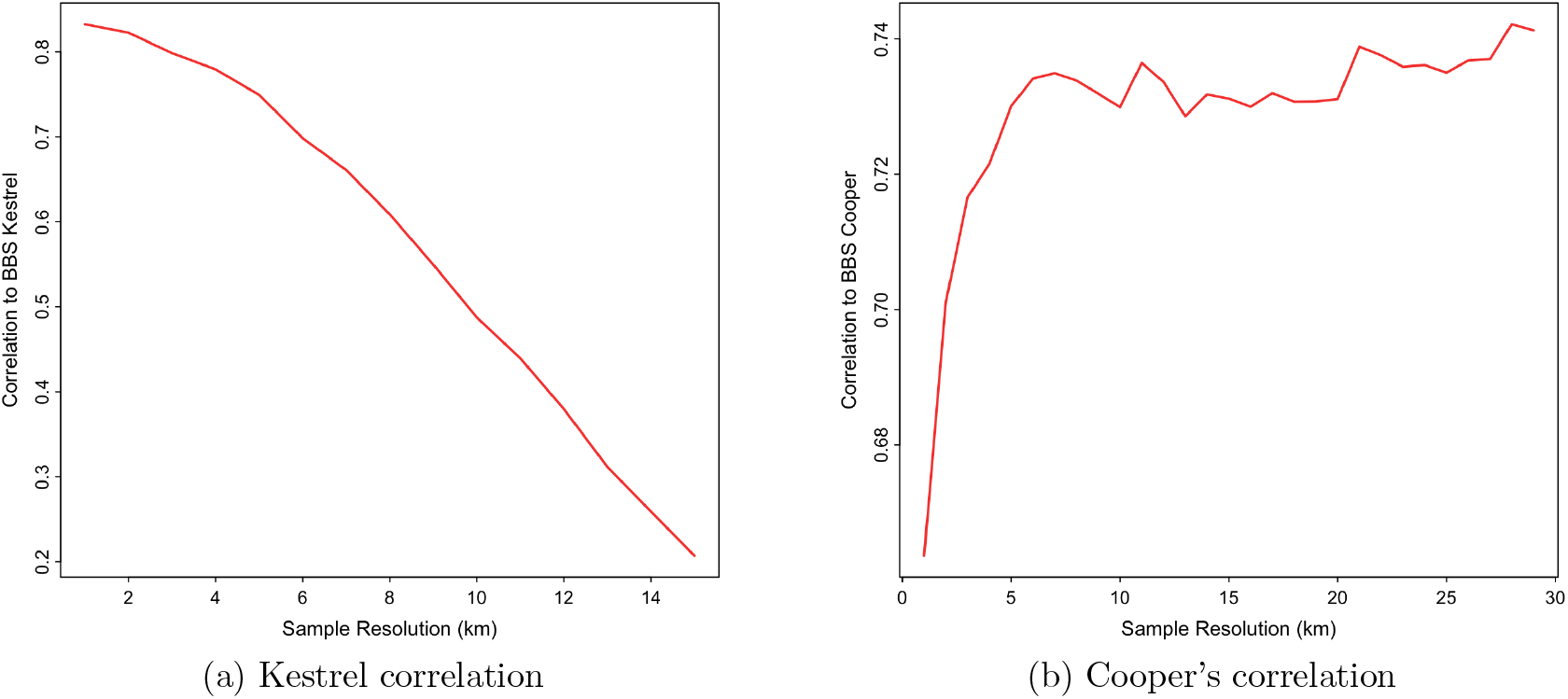
Comparison of subsampling resolutions

**Figure 7:**
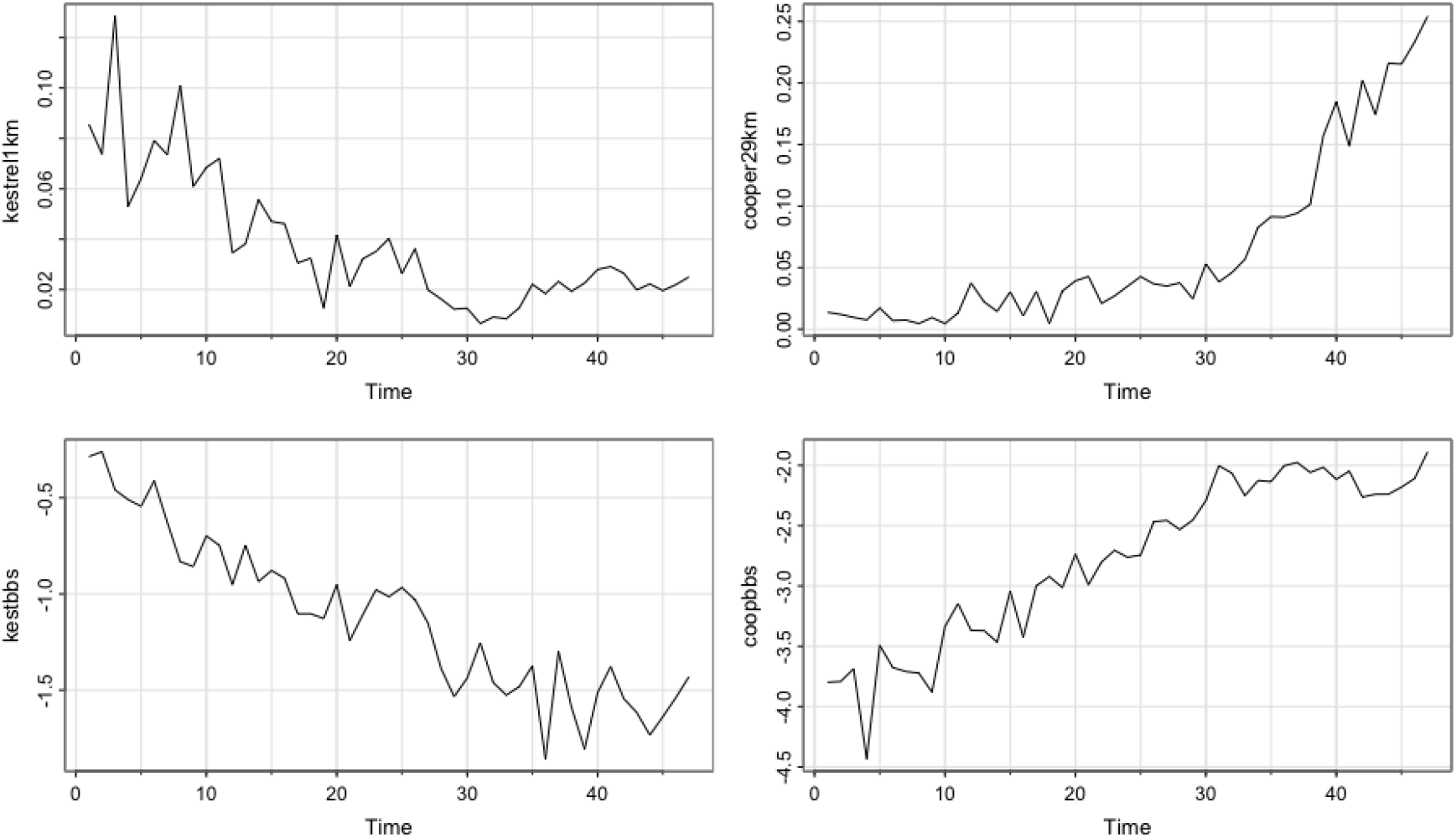
Series from subsampled checklists

**Figure 8:**
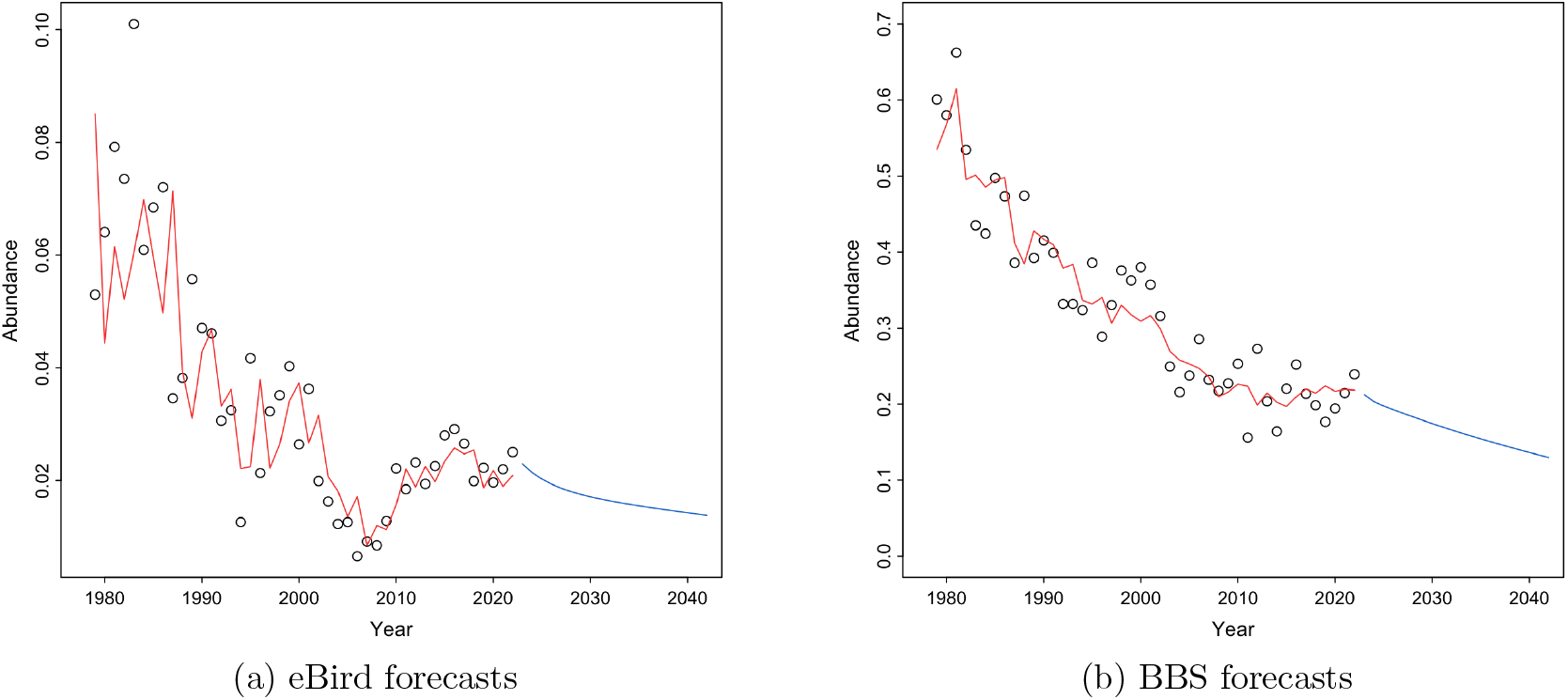
Comparison of forecast results for Kestrel abundance

### 2.3 Time Series Analysis

Autoregressive Distributed Lag (ARDL) models are fit to each of the sampled series. The model is Hanck et al. (2021)

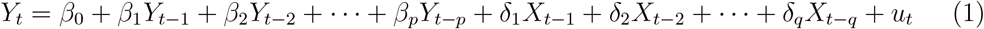

where *Y*_*t*_ is the target series at time *t*, and *X*_*t*_ is an external predictor at time *t. β*_0_ is the intercept, and *β*_*p*_ is the covariate coefficient of the *p*- th lag of the target series and *δ*_*q*_ is the covariate coefficient of the *q*-th lag of the external series. *β*_0_ is the intercept of the data, and *u*_*t*_ is an error term with mean 0 given all of the lags.

The most accurate model is selected based on the corrected Bayesian Information Criterion (BIC) based on the maximum likelihood estimation. Forecasts can then generated based on this model. BIC is generally the better criterion since there is a penalty for lag length in the log(*T*) term. BIC is calculated as

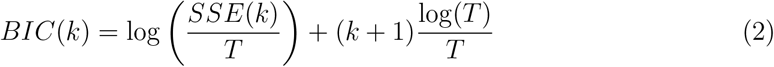

where *k* = *p* + *q*, the number of regressor parameters, *SSE*(*k*) is the sum of square errors, and *T* is the length of the time series. BIC attempts to find the true lag length, which is why it is the criterion used.

Then, multiple forecasts will be generated with different changes to the Cooper’s Hawk population regressor to determine which way Kestrel populations will trend.

Here, we describe the process of fitting the time series model to the subsampled eBird data. We provide details on model estimation, including the specific parameter settings and considerations for capturing seasonal and trend components. Using the fitted model, we make a forecast to predict future population trends, interpreting the results in the context of the dynamics of the American Kestrel population.

After subsampling, the amount of Kestrels and Cooper’s Hawks found each year was simply divided by the number of checklists in the sample each year to get an accurate description of the study region based on the verification in the simulation study.

In eBird, users have the option to report that a bird was present without mentioning the count. In this case, all observations with “X” were shifted to the median observation count of 1 for both Kestrels and Cooper’s Hawks.

To account for the possibility of negative forecasted abundance, the forecasts and model fitting are done based on the log-transformed data and exponentiated back to display the true forecast.

#### 2.3.1 Regression Results (BIC) and Forecast for Top 3 Models

The ARDL order is chosen by BIC for the BBS data to be (1, 2). The sampled series with eBird with the same order is chosen, and overall, show a significant overall negative effect based on the model coefficients, as both tables below show. In both cases, a non-statistically significant first lag is followed by a statistically significant second lag.

The forecast results for both series then provided by exponentiation of the log-transformed abundances and were highly similar, showing the validity of eBird as a long-term dataset.

## 3 Simulation Study

In this section, we present the details of our simulation study, including the creation of synthetic data to mimic both Breeding Bird Survey (BBS) and eBird data, the number of repetitions (over 200), and the confidence intervals of estimated parameters based on these repetitions. We discuss the implications of the findings from this study.

### 3.1 Simulation Setting

This subsection describes the setup of our simulation study. Key elements include the length of synthetic data, the dimensions of the spatial grid (e.g., a 10 × 10 grid), and the number of repetitions (200+) we performed. These settings are chosen to replicate typical conditions in citizen science datasets like BBS and eBird. The simulation setup is done by creating a 10×10 grid with a length of 47 years. To simulate real-world collection of eBird data, we increase the amount of ‘checklists’ that we are sampling over time. The ground truth model is constructed based on a reasonable estimation for the intercept and slope of Kestrels and Cooper’s Hawks. These trends are then tested through repeated simulation. This increasing checklist abundance estimates the amount of Kestrels found on checklists in a spatial region each year. This process is repeated 200 times to estimate the *α*_1_ and *β*_1_ parameters with an empirical confidence interval..

### 3.2 Ground Truth Model

Here, we outline the ground truth model assumptions for the synthetic abundance data of American Kestrels and Cooper’s Hawks. This model serves as the baseline against which we assess our framework’s performance in estimating abundance trends, simulating true population dynamics under predefined conditions.

This spatial randomness can be shown through a Gaussian process in a Gaussian Random Field (GRF). The GRF assigns a random value to an individual grid cell, but still follows a spatial dependence structure. The value which changes the intercept and slope is randomly determined by independent GRFs, denoted as *f*_*a*_(*s*) and demonstrated in Figure 9. The intercept and slope estimates, 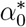 and 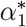 are given reasonable values based on estimated change over time and rarity on checklists. At a point on the grid *s*, the GRF values can add spatial-dependent randomness to both of these parameters mathematically as

**Figure 9:**
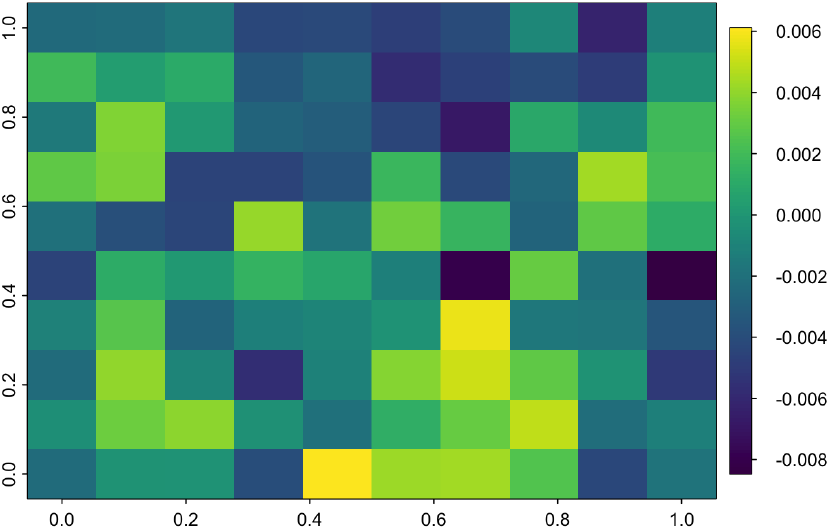
Gaussian Random Field on a 10×10 grid

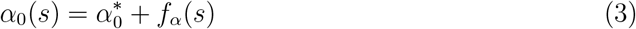

and the same for 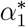. Assuming the model is the same as described in Equation 1, we can describe each of the abundances in a spatial grid at box *s* at time *t* as

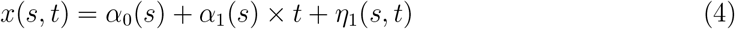

where *η* is N(0, 0.1). We can then model *x* as the series representing Cooper’s Hawks, and assume that *α*_1_*\** is positive to represent the increasing trend.

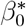 and 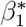 then represent the intercept and slope of the American Kestrel series. Following a similar process to the construction of the Cooper’s Hawk series in Equation 4 with GRFs being added to the parameter estimation as in Equation 3. Instead of making time the independent factor, we make the Cooper’s Hawk series the independent factor as in Equation 5 where *ϵ* is another normal error term.

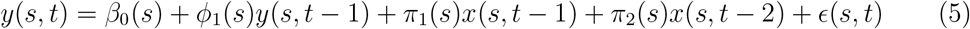

To take the region consensus, we take the average of the parameters at each spatial grid for *n* grid boxes using the operations in Equation 6, 7, and 8.

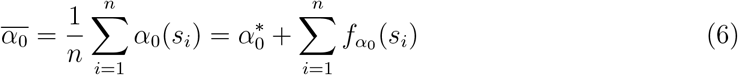

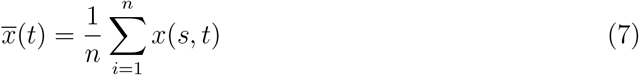

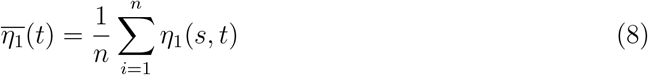

Thus, over the entire spatial grid,

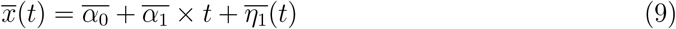

and

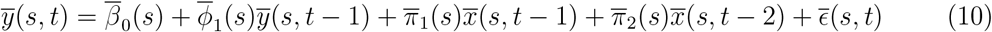

### 3.3 Distribution of Checklists

This subsection explains the methods used to introduce sampling bias into the synthetic data, designed to reflect the uneven geographical and temporal distribution of eBird observations. We describe how this sampling bias is embedded in the synthetic data to simulate real-world data collection tendencies.

eBird data contains uneven geographic and temporal distributions. Some geographic areas have more checklists than others, and the amount of checklists have increased over time. We then create a similar process in the synthetic dataset. Over the spatial grid, every 10 cells receives an increase in checklists to ‘sample’ from. The amount of checklists is also increasing over the sample period. The detection frequency is applied in a Poisson random variable with the amount of kestrels being selected from the ground truth.

### 3.4 Performance Measurement

In this subsection, we present the results of the simulation, showing how our estimators perform in relation to the ground truth values across 200+ simulation repetitions. We provide empirical 95% confidence intervals to demonstrate the accuracy and reliability of our estimators in capturing the true abundance trends.

## 4 Discussion

The results from the analysis and the forecasts supported the initial hypothesis that the quick rebound of predatory species, combined with the habitat changes by the Kestrel population after the ban of DDT, has continued to drive Kestrel numbers down. The analysis also supports the conclusion that, while the ban on DDT was helpful for conservation purposes, it may not be enough to save certain bird species such as the Kestrels due to other emerging factors.

In the samples of eBird data, at the representative 1 km and 29km samples of the data, we can see that the increasing Cooper’s Hawk abundance in the forecasts led to a decrease in Kestrel abundance, along with the external regressor showing a negative total coefficient.

### 4.1 Analysis of Subsampling and Simulation Results

The subsampling was able to alleviate spatiotemporal biases for both species at the 3 and 10 km resolutions as shown in Figure 7. At these resolutions, the series resembled the general BBS trend for their species. However, at larger resolutions, there were some limitations to the success of the sampling for kestrels.

First, the sampling process uses case control to oversample detections. Since Kestrels and Cooper’s Hawks are rare, oversampling detections addresses the class imbalance between detection and non-detection checklists. This oversampling worked well for 3 km and 10 km grid boxes, since most checklists in the grid box were representative of the grid box. However, when the boxes are larger, a spatiotemporal bias is reintroduced. As the number of checklists increased, the number of gridboxes that contained a checklist also increased. The new boxes bordered open areas because of their large size (30 km) and Kestrels are more frequent in these areas. Checklists outside of Kestrel habitat could be boxed into the large grid size and are not represented creating a bias in this sample. This spatiotemporal bias is illustrated in Figure 10.

**Figure 10:**
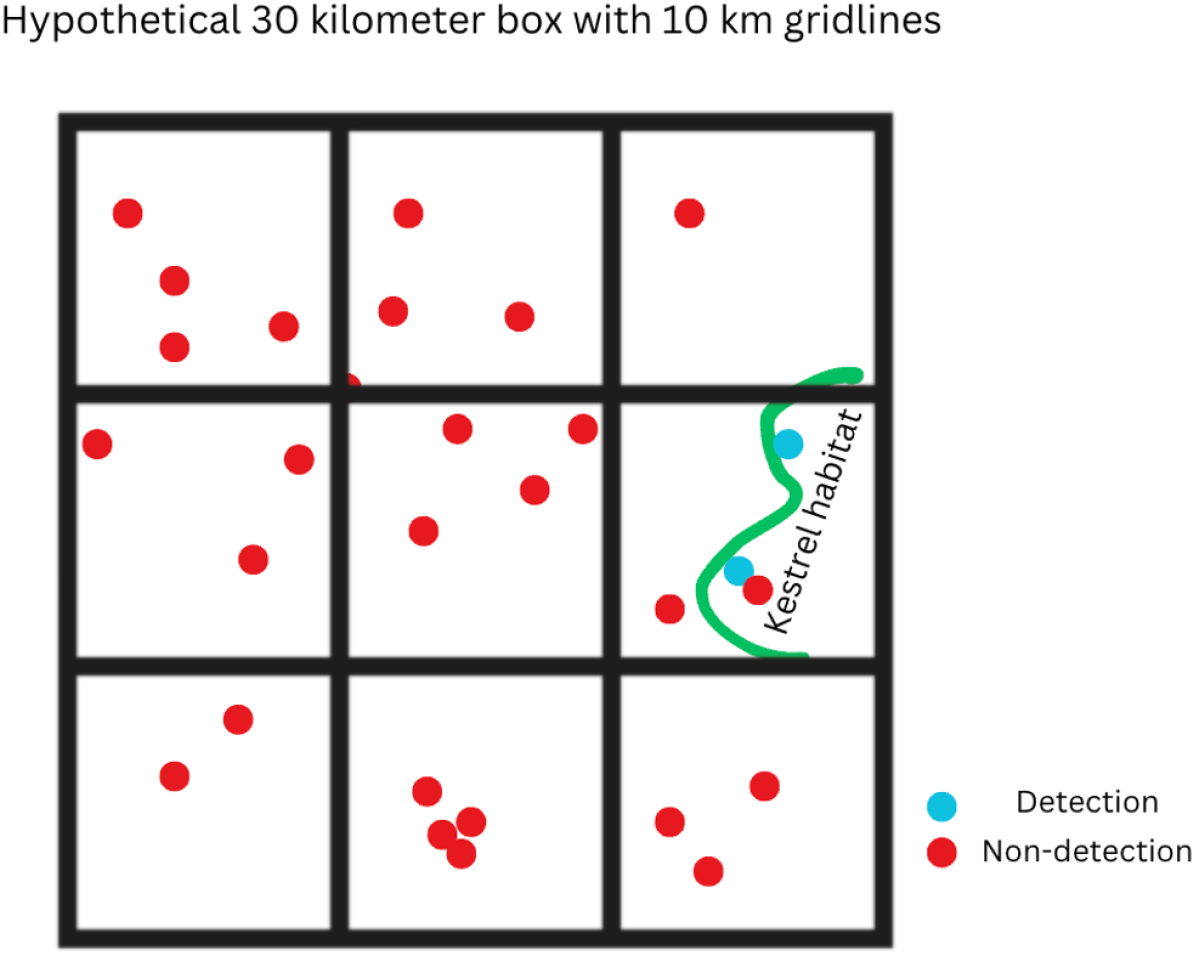
Case Control Sampling on a Hypothetical Grid

The simulation process had high accuracy at discerning the spatial inhomogeneity in the data. The simulated series’ spatial dependence on the GRF made spatial variation important to models at individual grid cells, showing that the reconstruction of the data was accurate as a spatial consensus.

### 4.2 Analysis of Regression & Forecasts

The BIC values in Table 1 and 2 allow us to assume forecast validity. Among similar datasets, this model performs the best in its ability to forecast. Both forecasts and spatial coefficients were similar and all of the forecasts in the representative samples pointed to the increase in Cooper’s Hawks pointing to decreases in Kestrel populations. The significant negative coefficients in the models show that there is a degree of truth to the hypothesis.

**Table 1:** BBS model.

| BBS | Coefficient (log) | Std. Error | t value | $Pr(> \vert t\vert )$ |
| --- | --- | --- | --- | --- |
| Intercept | -2.09807 | 0.37195 | -5.641 | 1.51e-06 |
| Kestrel 1 lag | 0.21102 | 0.13467 | 1.567 | 0.12502 |
| Cooper 1 lag | -0.13306 | 0.09029 | -1.474 | 0.14840 |
| Cooper 2 lags | -0.28459 | 0.09499 | -2.996 | 0.00468 |

**Table 2:** eBird model.

| eBird | | Coefficient (log) | t value | $Pr(> \vert t\vert )$ |
| --- | --- | --- | --- | --- |
| (Intercept) | -1.10963 | 0.53030 | -2.092 | 0.0428 |
| Kestrel 1 lag | 0.74564 | 0.10470 | 7.122 | 1.27e-08 |
| Cooper 1 lag | 0.16573 | 0.09703 | 1.708 | 0.0954 |
| Cooper 2 lags | -0.21387 | 0.09212 | -2.322 | 0.0254 |

### 4.3 Research Novelty and Potential for Further Exploration

Past research has focused on standard/traditional linear models to observe species trends. With the amount of data increasing exponentially, especially large, crowd-sourced citizen science datasets (such as eBird), analyzing the data through time-series analysis methods, while automatically accounting for biases and errors, is a novel addition that opens up many new possibilities. These methods can be used to both explain ecological relationships, as well as identify new dependent and independent variables that affect the environment, in this particular case, the real drivers impacting the decline in Kestrel population. Additionally, this research is focused on building models that can be enhanced, reused and built upon to understand impact on a broader scale.

Further research could focus on the inclusion of more covariates into the ARDL regression, and that time-based models with more parameters such as General Autoregressive Conditional Heteroskedasticity (GARCH) models can aid in this analysis. Machine learning can also be added to this analysis, and Automated ML could build models quicker and efficiently. Further enhancements of the statistical analysis present can reshape understanding of our bird populations and subsequently determine impactful steps for their conversation.

## Acknowledgements

We would like to thank Dr. Jim Bednarz for his insights on American Kestrels, and the University of North Texas for providing the platform to conduct this research on campus.

